# Carrier based formulation of plant growth promoting *Rhizobium* species and their effect on *albizia procera* seedling

**DOI:** 10.1101/2022.12.15.520691

**Authors:** Neerja Rana, Arti Ghabru, Meenakshi

## Abstract

The carrier-based formulation of rhizobial cultures in general, excels to register their effects on various plant growth parameters of *Albizia procera* seedlings. Carrier based bio-inoculant prepared with rhizobial isolate FA6 of Uttrakhand performed better for all plant growth characteristics. Charcoal based application of rhizobial isolates also increased the N, P, K content of soil. In case of seedlings treated with charcoal-based formulation of rhizobial isolate from Himachal Pradesh maximum available N (323.30 kg/ha), P (38.97 kg/ha) and K (395.9 kg/ha) content was recorded in T_2_ (Charcoal based formulation of BA2 strain+20kg/ha N) which was significantly superior to T_3_ and uninoculated control. Among the charcoal-based formulation of rhizobial isolate from Uttrakhand seedlings under treatment T_2_ (Charcoal based formulation of FA6 strain+20kg/ha N) registered maximum available N (327.7kg/ha), P (39.01 kg/ha) and K (397.4 kg/ha) content in soil. Charcoal based application of rhizobial isolates significantly influenced the activity of soil enzymes (Dehydrogenase and Phosphatase). Application of Carrier based formulation of rhizobial isolate BA2 from Himachal Pradesh showed maximum dehydrogenase (11.20 μg TPF g^-1^h^-1^) and phosphatase (536.2 μmolL^-1^g^-1^h^-1^) activity as compare to uninoculated control. Whereas, among the rhizobial isolates from Uttrakhand maximum dehydrogenase (11.39 μg TPFg^-1^h^-1^) and phosphatase (545.4 μmol L^-1^g^-1^h^-1^) activity was also recorded with T_2_ (charcoal-based formulation of strain FA6+20kg/ha N) which was significantly superior to T_3_ and uninoculated control.

## Introduction

It has been well-known fact that the inoculation of legumes with bacteria is moderately essential for their suitable establishment when grown in new place (Afzal and Bano,2008). Rhizobium are soil microorganism that either saprophytes (live on plant residues) or endophytes (entirely within plants) or in close connotation with the plant roots. They play a dominant role in the Nitrogen supply of most soil ecosystems through their ability to fix Nitrogen in symbiosis with legumes.

The agricultural benefits possible from use of selected, high-Nitrogen fixing strains of rhizobia can be realized only when farmers obtain and properly use high-quality inoculants on their legume seeds or soil before planting. Technology on growing rhizobia, preparing inoculants with suitable carrier materials and distributing viable inoculants to farmers is essential. But the key constraint in successful commercialization of bacterial inoculant is overcoming difficulties in formulating a viable, cost effective and user friendly final product. Development of formulations with increased shelf life and broad spectrum of action with consistent performance under field conditions could be beneficial for commercialization of the technology at a faster rate. The usage of carrier materials for the microbial inoculants proves to be beneficial to protect the bacteria and have long been practiced (Ardakani et *al*., 2010; Fuentes-Ramirez and Caballero-Mellado, 2005). Carrier material provides a medium on which micro organisms multiply to a reasonably high number with long shelf life and it is one of the basic considerations for bio-inoculants production. The benefits of incorporating microorganisms in carrier material are: easy-handling, long-term storage and high effectiveness of bioinoculant. The carriers for bioinoculant preparation include soil (John, 1966), filter mud, lignite (Philpotts, 1976), compost of coir dust, charcoal, manure, compost powdered coconut shells, ground teak leaves (Tilak and Subba Rao, 1978) coal (Crawford and Berryhill, 1983) and talc powder (Vidhaysekaran *et al*., 1997). Also, the combinations of these substances have been tried across the world by various researchers. Chao and Alexander (1983) studied mineral soil as carrier for rhizobia inoculant and reported that survival of bacterium inoculants applied to seed at lower temperature was equal to that of peat based culture.

Pahari *et al*. (2017) evaluated the plant growth promoting activity of charcoal and talc-based formulation of *Bacillus* species on mung bean and rice. They reported that charcoal-based formulation depicts higher plant growth promoting activity in comparison with talc-based formulation. They further reported that bio-inoculants increased organic carbon, nitrogen, phosphorous and potassium in soil, there by promoting growth of mung bean and rice in respect to control. To the best of our knowledge no work has been done on carrier based bioinoculant preparation from/for rhizobia isolated from tree legumes.

## MATERIALS AND METHODS

### SAMPLE COLLECTION, inoculum preparation, and inoculation

Pods and rhizospheric soil samples were collected from 15 to 25 year old matured and healthy *Albizia procera* (Roxb.) trees from different geographic locations of Himachal Pradesh and Uttarakhand (Table 1). In each location three representative sites were selected and three plant samples were collected from each site. The samples were placed in plastic bags and stored in Laboratory for further isolation and analysis work. Sowing of overnight soaked pod-less seeds was done, and a nursery trial was set up. Isolation and enumeration of bacteria associated with the rhizospheric region of *Albizia procera* soil samples (collected from geographic locations of Himachal Pradesh and Uttrakhand) were done. The isolated rhizospheric colonies were prepared as microbial inoculants and were treated (soil drenching) on the same seed sown poly bags as the geographic area after 15 days for the establishment of the rhizospheric niche. The soil drenching with inoculum was carried out till the end of the nursery trial.

**Table 1.** Geographical locations and habitat data of various seed sources of *Albizia procera* in North western Himalayan Region.

| Location | District | Altitude<br>(amsl) | Latitude | Longitude | Rain<br>fall<br>(mm-<br>/yr-2016) | Plant<br>height<br>(m) | Plant<br>diameter<br>(cm) |
| --- | --- | --- | --- | --- | --- | --- | --- |
| UHF Nauni | Solan | 1265.1 | 30°53'58.46"<br>N | 76°49'31.81"<br>E | 1612 | 20.5 | 24.6 |
| Baddi | Solan | 433 | 30°53'58.46"<br>N | 76°49'31.81"<br>E | 1035 | 26.4 | 30.3 |
| RHR&TS<br>Jachh | Kangra | 407.3 | 32°9'54.31"<br>N | 75°30'<br>47.89"E | 1196 | 22.7 | 27.2 |
| Gaggal | Kangra | 752.2 | 32°<br>9'36.75"N | 76°15'<br>42.67"E | 1831 | 22.2 | 26.5 |
| RHR&TS<br>Dhaulakuan | Sirmour | 534 | 30°27'12.27"<br>N | 77°28'44.46"<br>E | 1742 | 23.6 | 28.3 |
| FRI<br>Dehradun | Dehradun | 671 | 30°<br>20'38.89"N | 77°<br>59'50.19"E | 1351.6 | 25.1 | 31.8 |
| ITBP Campus | Dehradun | 626 | 30°18'18.07"<br>N | 78° 0' 1.27"E | 1787.2 | 20.8 | 29.1 |
| Pant Nagar<br>University | Udham<br>Singh<br>Nagar | 229 | 29°0'29.30"<br>N | 79°29'16.69"<br>E | 1351.6 | 25.4 | 29.4 |
| Ram Nagar | Nainital | 558 | 30°10'34.81"<br>N | 78°13'5.49"E | 734.4 | 20.5 | 23.8 |
| Lal Kuan<br>Forest Range | Nainital | 261 | 29°3'<br>37.34"N | 79°31'19.06"<br>E | 1104.2 | 24.3 | 28.2 |

#### Isolation of rhizobia from nodules

The rhizobia isolation was performed from the nodules of root *Albizia procera* (six-month-old) seedlings grown under the net house environments. The poly bags containing seedlings from each site with the best nodulation was selected for uprooting. Seedling roots were washed thoroughly to remove soil till the nodule became visible clearly. Nodules were then separated in an autoclaved petri dish containing 0.1% HgCl_2_ for a few seconds (20 to 30 seconds) and washed thoroughly in autoclaved distilled water several times. A sterilized Forcep or needle is used to puncture the nodules for a whitish or milky bacteroid suspension. The suspended bacterial culture is then streaked on a specific medium of yeast extract mannitol agar plate incubated for 3-5 days at 28±2°C.

#### Evaluation of bacterial nodules

Authentication test of isolates from nodules was done as prescreening of *Rhizobium* obtained from nursery trial. Following tests were done to evaluate the presence of *Rhizobium*.

#### Congo red test

An aliquot of 2.5 ml of 1 percent solution of the congo red dye in water was added to a liter of YEMA medium. All the purified rhizobial isolates were streaked on CRYEMA medium and incubated for 3-5 days at 28±2°C. Rhizobial colonies were observed for absorption of congo red dye (Vincent, 1970).

#### Bromothymol blue test

The YEMA medium containing bromothymol blue was streaked with isolated strains and incubated for 3-5 days at 28±2°C. Rhizobial colonies were observed either for yellow color due to production of acids or blue color due to production of alkali (Norris, 1965).

#### Hofer’s alkaline test

Yeast extract mannitol broth (pH 11.0) in a test tube was inoculated with a loopful culture and incubated for 2-5 days at 28±2°C (Hofer, 1935). This test is based on the fact that *Agrobacterium* grows at a higher pH level, whereas *Rhizobium* is unable to grow at pH 11.0.

#### Ketolactose agar test

Ketolactose agar plates were streaked with isolated microbes. After incubation for 4-6 days at 28±2°C, the plates were flooded with Benedict’s solution. The composition of Benedict’s solution consists of two solutions: Solution A (Sodium citrate-173 g, Anhydrous sodium carbonate-100 g, distilled water-600 ml) and Solution B (Copper sulphate-17.3 g, distilled water-100 ml). The solutions A and B were prepared separately, and later solution B was mixed with solution A and then filtered. The resultant solution was cleared and had a transparent blue color. After pouring Benedict’s solution into the plates mentioned above, the excess of the solution was drained off, and plates were incubated for 1 h at 28±2°C. This test is based on the fact that *Agrobacterium* utilizes lactose by the action of enzyme ketolactase and shows yellow coloration around the bacterial colonies, whereas rhizobia cannot utilize the lactose (Bernaerts and Deley, 1963).

#### Plant infection assessment

Various isolates were assessed for their capability to nodulate *Albizia procera* plants grown in pots (plastic). All isolates of *Rhizobium* were grown up to 6 days at 28±2°C on YEM broth. *D. sissoo* seeds were inoculated with *Rhizobium* isolates (soaking seeds with 100 ml bacterial suspension) and sown in pots. After 75 days, plants were cautiously uprooted and observed for nodulation.

#### PGPR Test

PGPR traits of the nodular bacteria were observed. P-solubilization liquid assay and plate assay (P-solubilization solubilization efficiency), siderophore liquid assay, and IAA production were performed. After authentication and evaluation of PGPR isolates, the selected isolates were molecularly characterized.

#### Quantitative assessment of phosphate solubilization

Medium (50 ml) was dispensed in Erlenmeyer flask OF 250 ml containing 0.5 percent tri-calcium phosphate (TCP) and autoclaved at 15 lbs per square inch of pure steam for 20 min. The flasks were inoculated with 1 percent (5 ml) of the 24 hold rhizobial suspension (OD 1.0 at 540 nm) and incubated at 28±2°C on a rotating shaker at 100 rpm for 72 h. The content was centrifuged at 15,000 rpm for 20 min at 4°C. The culture supernatant was used for the determination of the soluble phosphorous as described by Bray and Kurtz (1945).

#### Qualitative estimation of phosphate solubilization

Each purified isolate was seeded in a straight line on PVK medium as described by Pikovskaya (1948) and was incubated for 72 h at 28±2°C. The rhizobial solubilization of phosphorus was exhibited with clear zones produced around the isolated rhizobial colony. Colonies showing solubilization halos (>0.1 mm diameter) were selected. Percent solubilization efficiency was calculated as:SE(%)=Z−CC×100 Where,SE(%)=Solubilizationefficiency(%)Z=Cleardiameter(inmm)C=Diameterofcolony(inmm) The diameter of the halo zone was calculated by deducting the colony size from the total size. PSI (Phosphate solubilization index) was calculated by using the formula (Edi-Premono *et al*., 1996).

#### Quantitative estimation of siderophore

Siderophore estimation was done using CAS (Chrome-Azurol S) assay as described by Schwyn and Neilands (1987). Cell-free extract (0.1 ml) of culture supernatant along with CAS (Chrome-Azurol S) assay solution (0.5 ml) mixed with the 10 μl 0.2M 5-Sulfosalicyclic acid solution (shuttle solution). And then, left at room temperature (RT) for 10 minutes, and at 630 nm, absorbance was noted. The reference (r) sample was prepared by using the same constituents except for the cell-free extract of the culture supernatant. On the other side, a minimal medium was used as a blank. And finally, the siderophore unit was calculated by:%Siderophoreunit=Ar−AsAr×100

Where,As=Testsampleabsorbanceat630nmAr= References ampleabsorbanceat630nm

#### IAA quantitative estimation

The method described by Gordon and Palleg (1957) was used to determine IAA. Rhizobial cultures were grown in Luria Bertani broth (amended with 5 mM L-tryptophan, 0.065 sodium dodecyl sulfate, and 1% glycerol) for 72 h at 28±2°C under shake conditions. The cultures were centrifuged at 15,000 rpm for 20 min, and the supernatant was collected. 3 ml of supernatant was added to 2 ml Salkowski reagent (2 ml 0.5 M FeCl_3_□+□98 ml 35% HClO_4_). The tubes containing the mixture were left for 30 min (in the dark) for the development of pink color. The intensity of color was measured spectrophotometrically at 535 nm. The concentration of IAA was estimated by preparing a calibration curve using IAA as standard (10-100 μg/ml).

#### Sequence and phylogenetic analysis

Representative rhizobial isolates of each genotypic profile were chosen for 16S rDNA partial sequencing. The sequencing was performed from Eurofins lab, Bangalore, India, using both forward and reverse primers, as mentioned in section 3.9.3. The sequence alignment was done using the Clustal W and manually edited using the Bioedit package (Thompson *et al*., 1994; Hall, 1999). The cladograms were constructed by using the neighbor-joining (NBJ) method (Saitou and Nei, 1987) with the Kimura-2-parameter model (Kimura, 1980) and were bootstrapped using the software programs in the MEGA 6.0 package (Tamura *et al*., 2013)

#### Molecular characterization

Representative rhizobial isolates of each genotypic profile were chosen for 16S rDNA partial sequencing. The sequencing was performed from Eurofins lab, Bangalore, India, using both forward and reverse primers, i.e., universal primers for 16S rDNA amplification of rhizobial isolates use4d were pA (5’-AGAGTTTGATCCTGGCTCAG-3’) and pH (5’-AAGGAGGTGATCCAGCCGCA-3’) designed by Edwards et al. (1989).

The sequence alignment was done using the Clustal W (Thompson et al., 1994) and manually edited using the Bioedit package (Hall, 1999). The cladograms were constructed by the neighbor-joining method (Saitou and Nei, 1987) with the Kimura-2-parameter model (Kimura, 1980) and were bootstrapped using the software programs in the MEGA 6.0 package (Tamura et al., 2013).

### PREPARATION OF LIQUID RHIZOBIAL FORMULATION

#### Seed sterilization, Preparation of liquid formulation and Seed inoculation

Seeds of *Albizia procera* were surface sterilized in 0.2 per cent mercuric chloride (Hg Cl_2_) solution for 2-3 min and rinsed at least 6-8 times with sterilized distilled water. Sterilization was checked by placing surface sterilized and unsterilized seeds on nutrient agar plates. The bacterial growth, if any, around the seeds was recorded after 48 h of incubation at 28±2°C. The population density (1.5 O.D. at 540 nm) that resulted in formation of 10^8^ cfu/ml of rhizobial isolate was used for preparation of liquid formulation. Seeds were soaked in liquid culture having 10^8^cfu/ml for duration of 3 to 4 h in sterilized petriplate and thereafter the seeds were sown.

### SELECTION AND EVALUATION OF CARRIER MATERIAL FOR SURVIVAL OF SELECTED PGPR

Three different types of carrier materials viz., charcoal, vermicompost and vermiculite were used to study the shelf life of *Rhizoium*.

The bacterial suspension resulted in population density (1.0 O.D at 540 nm) and containing 10^8^ cfu/ml cells of rhizobial isolates were added to 100 g charcoal, 100 g of dry vermiculite,100g of vermicompost and 35 per cent moisture content was adjusted. A suitable carrier in ratio of 1:1 (w/v) was used. The amount of inoculum broth was below carrier moisture holding capacity to make mixture friable in texture. The carrier was autoclaved in flasks before inoculum was added. The desirable amount of liquid suspension was added to the carrier. (Somasegaram and Hoven.,1985). The number of viable count (cfu/g) was taken after 6^th^ day incubation and presented as counts on 0^th^ day. Count was taken after 2, 4 and 6 months of storage of inoculants. The viable count was determined by following the standard serial dilution and plate count method.

#### Treatments

T_1_ = Vermicompost + BA2 rhizobial strain, T_2_ = Charcoal + BA2 rhizobial strain,T_3_ = Vermiculite + BA2 rhizobial strain, T_4_ = Vermicompost + JA4 rhizobial strain, T_5_ = Charcoal+JA4 rhizobial strain,T_6_ = Vermiculite + JA4 rhizobial strain, T_7_ = Vermicompost + FA6 rhizobial strain, T_8_ = Charcoal + FA6 rhizobial strain, T_9_ = Vermiculite + FA6 rhizobial strain, T_10_ = Vermicompost + IA5 rhizobial strain, T_11_ = Charcoal + IA5 rhizobial strain, T_12_ = Vermiculite + IA5 rhizobial strain

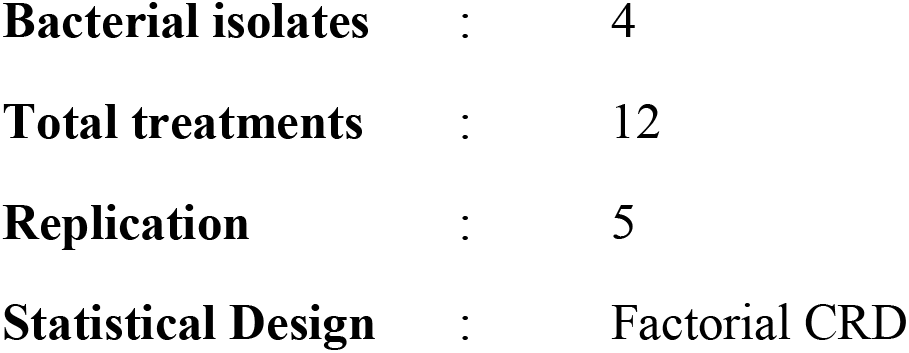

#### Observation

The growth of rhizobia in terms of log_10_ cfu/ml was recorded after 0^th^ day, 2, 4 and 6 months.

#### Evaluation of carrier based bioinoculant and chemical fertilizer on the growth and nitrogen fixation of selected seed source of *Albizia procera*

Effect of selected carrier based bioinoculant was evaluated on the growth and nitrogen fixation of selected seed source of *Albizia procera* under net house conditions.

#### Seed Treatment with Carrier Based Bioinoculant

##### Surface Seed sterilization

Seeds were surface sterilized in a 0.2 per cent mercuric chloride solution for two minutes and rinsed several times with sterilized distilled water to remove the contents of mercuric chloride.

##### Adhesive

The adhesive (sticker) used in this study was 40 per cent solution of jaggery (Gur). The jaggery solution was prepared by adding granular jaggery in small (1-2 g) lots under continuous stirring to 100 ml of distilled water heated to near boiling.

##### Seed inoculation with carrier based rhizobial inoculant

*Albizia procera* seeds were placed in sterilized flask (250 ml) and 10 ml of adhesive were added the flask was gently shaken for at least 60 seconds to coat all the seeds thoroughly with adhesive so that seeds appeared wet. Then 10 g of carrier based inoculants of rhizobia, was added to the flask. The flask was gently shaken until the seeds get coated with carrier based inoculant. Shaking was stopped and the seeds were immediately transferred to the sterilized Petri plate, and allowed to air dry.

##### Sowing of seeds

Five seeds treated with carrier based bioinoculant were sown at equidistance atleast 3 cm depth in each pot. After 15 days of seedling emergence, thinning was done and only one plant per pot was maintained.

##### Soil application of carrier based culture

Soil application of carrier based rhizobial culture was also done. For the soil application, 10g of carrier based rhizobial culture was mixed with soil in each pot before sowing of seeds.

#### Treatments

##### With rhizobial isolates from Himachal Pradesh

T_1_= Control,T_2_ **=** Charcoal based bioinoculant with rhizobial strain BA2, T_3_ = Charcoal based bioinoculant with rhizobial strain JA4

##### With rhizobial isolates from Uttrakhand

T_1_= Control, T_2_ **=** Charcoal based bioinoculant with rhizobial strain FA6, T_3_ **=** Charcoal based bioinoculant with rhizobial strain IA5

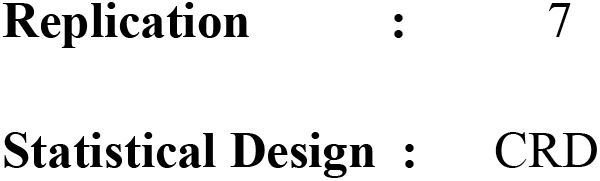

Hence, 1^st^ nursery comprises of maintaining the *D. sissoo* seedlings under a prominent environment. Morphological characters of seedlings were observed during the end of 1^st^ nursery trial.

Based on PGPR characters of rhizobia from 1^st^ nursery, effective rhizobia were added directly or along with nitrogen fertilizer in 2^nd^ nursery, and effects were observed on seedlings. Morphological characters of seedlings and the effect of PGPR(s) were observed (nitrogen-fixing potential or Nitrogenase activity) using Acetylene-ethylene assay during the end of 2^nd^ nursery trial.

##### Morphological evaluation

The shoot, root and, nodulation characteristics of *D. sissoo* seedlings were recorded after 6 months from the date of sowing for both 1^st^ nurseries.

##### Shoot characteristics

Shoot length: With the help of a meter scale, the length was measured from soil level to shoot the tip of each seedling.

The number of leaves/plant: A total leaf of each plant was counted except juvenile and very young leaves.

Area of the leaf (cm^2^): Area of the leaf was calculated by using leaf area meter LICOR LI-3000. Collar diameter (mm): By using Digital Vernier Caliper at the collar region, and the mean of three values was recorded. Collar diameter was expressed in millimeters (mm).

Shoot dry weight: Replication wise from every single treatment shoot dry weight was calculated and expressed in gram (g).

##### Root characteristics

Root length: Taproot length was measured by using a meter-scale for each seedling and expressed in centimeters (cm).

Root dry weight: Replication wise from every single treatment, root dry weight was calculated and expressed in gram (g).

##### Nitrogen fixation potential

The number of nodules: Nodule number of individual plants.

Nodule fresh and dry weight: Replication wise from each treatment nodule, fresh and dry weight was recorded. It was expressed in gram (g).

The Nodular Nitrogenase activity of selected *Albizia Procera* pot seedlings (T1 to T10) was determined using acetylene-ethylene assay (ARA) as described by Hardy et al. (1968). The following formula was used to calculate acetylene reduction:μmolesC2H2reducedg−1freshweighth−1=Peakheight(mm)×gasphase(ml)/weightofnod ule(g)×C2H2factor×timeofincubation(h)

##### Soil Enzyme (Phosphatase, Dehydrogenase)

The phosphtase enzyme estimation was carried out by method given by Tabatabai and Bremner (1969). The standard curve was prepared by p-nitrophenol (10-100 ppm). The result was expressed as μ mole of p-nitrophenol released per gram soil per hour (μmole p-nitrophenol g^-1^soil h ^-1^).

The dehydrogenase enzyme estimation was carried out by method given by Casida *et*.*al*. (1964). Extrapolate Triphenyl formazen (TPF) formed from the standard curve drawn in the range of 10 to 90 mg TPF ml^-1^ and results expressed as mg TPF formed per h per g soil.

##### Physicochemical, available nutrient and microbiological properties of soil mixture

Freshly prepared potting mixture was analyzed for important physico-chemical, available nutrient status and microbiological properties before and at the end of the experiment by adopting the following standard procedures:

##### pH and electrical conductivity

The soil pH was determined in 1:2.5 (soil: water suspension) and the electrical conductivity of the supernatant liquid was recorded and expressed in dSm^-1^ (Jackson, 1973).

##### Organic carbon

Organic carbon was determined by Chromic acid titration method of Walkley and Black (1934).

##### Available nitrogen

Available nitrogen was determined by alkaline permagnate method of Subbiah and Asija (1956).

##### Available phosphorous

0.5 N Sodium Bicarbonate (NaHCO_3_) at 8.5 pH was used to extract available phosphorus (Olsen *et al*., 1954) and determined spectrophotometrically.

##### Available potassium

Available potassium was extracted by normal neutral ammonium acetate (Merwin and Peech, 1951) and determined on flame photometer.

##### Viable microbial count

The soil was analyzed for viable microbial count by taking 1 g of soil in 9 ml sterilized water blank and the soil suspension was diluted in 10 fold series, then microbial count was determined by standard pour plate technique on different media as described by Subba Rao (1999). The count was expressed as colony forming units per gram of soil (cfu/g soil).

##### Statistical Analysis

The data recorded on plant, soil and microbiological properties were statistically analysed by using MS-Excel and OPSTAT packages (Sheoran *et al*., 1998). The mean values of data were used for the analysis of variance (ANOVA) as described by Panse and Sukhatme (2000) by using Complete Randomized Design as given in Table 2.

**Table 2.**
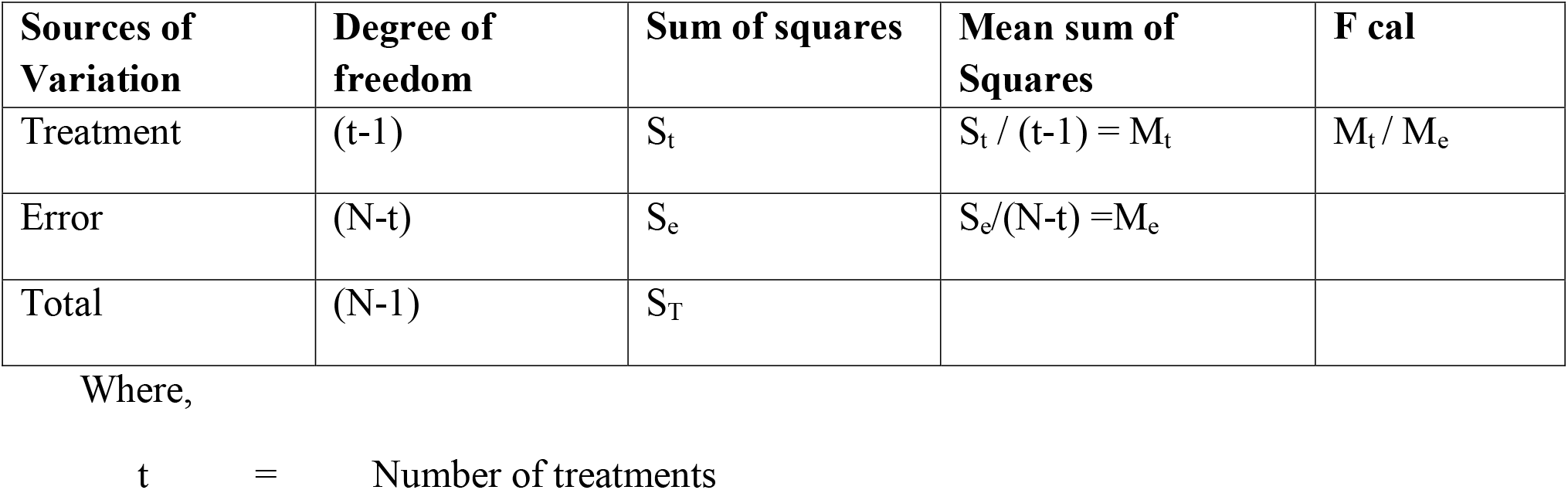
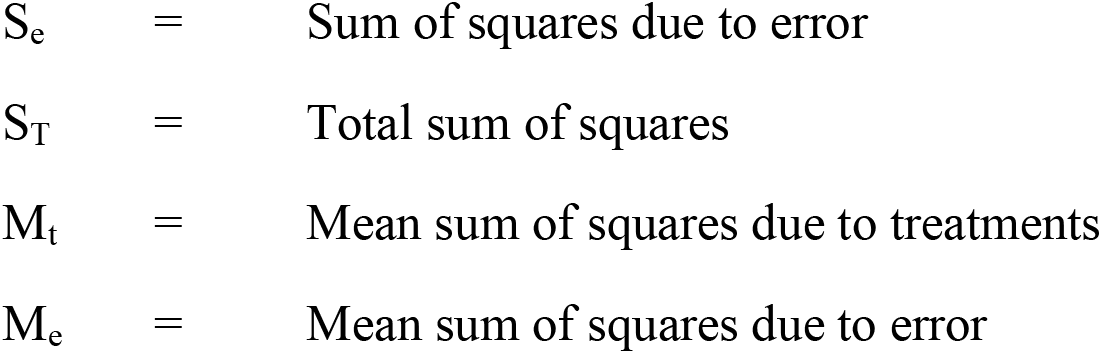
Analysis of variance (ANOVA) by Completely Randomized Design (CRD)

| Sources of Variation | Degree of freedom | Sum of squares | Mean sum of Squares | F cal |
| --- | --- | --- | --- | --- |
| Treatment | (t-1) | $S_t$ | $S_t / (t-1) = M_t$ | $M_t / M_e$ |
| Error | (N-t) | $S_e$ | $S_e / (N-t) = M_e$ | |
| Total | (N-1) | $S_T$ | | |

|  |  |  |
| --- | --- | --- |
| $S_e$ | = | Sum of squares due to error |
| $S_T$ | = | Total sum of squares |
| $M_t$ | = | Mean sum of squares due to treatments |
| $M_e$ | = | Mean sum of squares due to error |

The ‘F’ calculated value was compared with ‘F’ table value at 5% level of significance. If ‘F’calculated value is higher than ‘F’ table value, the treatment effect was considered significant i.e. one of the treatment pair differ significantly and critical difference (CD) was calculated.

The standard error of mean SE(m) and critical difference (CD) for comparing the means of any two treatments was calculated as below:

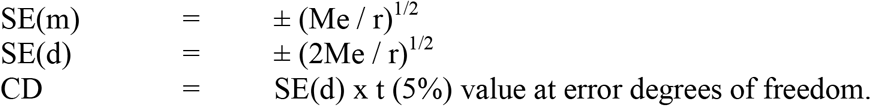

Where,

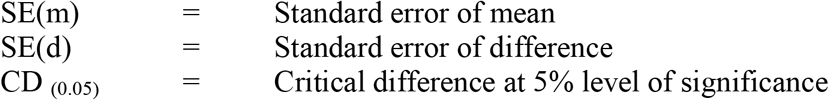

## RESULTS AND DISCUSSION

The present investigation was conducted under net house condition at the Department of Basic sciences, Dr. Yashwant Singh Parmar University of Horticulture and Forestry, Nauni.

### Isolation of rhizobia from root nodules of *albizia procera* seedlings grown under net house

The isolation of rhizobia was done from root nodules of six months old *Albizia procera* seedlings grown in net house. The growth on the YEMA medium was counted and reported as cfu**/**g of root. A significant variation in the rhizobial population colonizing the roots of *Albizia procera* seedlings was recorded and presented in Table 3.

**Table 3.** Isolation of rhizobial strains from root nodules of *Albizia procera* seedlings grown under net house.

| | Sites of sample collection | Isolate name | No. of isolate | Rhizobial Population<br>($\times 10^2$ cfu/g root) |
| --- | --- | --- | --- | --- |
| **Himachal Pradesh** | UHF Nauni | N A (Nauni *Albizia*) | 8 | 76.3 |
|  | Baddi | BA (Baddi *Albizia*) | 11 | 84.1 |
|  | RHR&TS Jachh | JA (Jachh *Albizia*) | 6 | 80.1 |
|  | Gaggal | GA (Gaggal *Albizia*) | 7 | 71.8 |
|  | RHR&TS Dhaulakuan | DA (Dhaulakuan *Albizia*) | 6 | 73.6 |
|  |  |  |  | CD<sub>(0.05)</sub><br>3.86 |
| **Uttarakhand** | FRI | FA (FRI *Albizia*) | 8 | 85.8 |
|  | ITBP | IA (ITBP *Albizia*) | 5 | 79.0 |
|  | Ram Nagar | RA (Ram Nagar *Albizia*) | 4 | 83.4 |
|  | Pant Nagar | PA (Pant Nagar *Albizia*) | 6 | 77.3 |
|  | Lal Kuan | LA (Lal Kuan *Albizia*) | 5 | 70.6 |
| CD (0.05) |  |  |  | 2.48 |

Among 5 seed sources of Himachal Pradesh maximum (84.1×10^2^ cfu/g root) rhizobial count on YEMA medium was noted from Baddi seed source whereas Gaggal seed source showed the minimum (71.8×10^2^ cfu/g root) rhizobial count. In case of 5 seed sources among Uttrakhand maximum (85.8×10^2^ cfu/g root) rhizobial count on YEMA medium was noted from FRI seed source whereas Lal Kuan seed source showed the minimum (70.6×10^2^ cfu/g root) rhizobial count. The variation in rhizobial population may be ascribed to influence of root exudates, age of plant, bioinoculant and physico-chemical properties of soil and environmental conditions (Wieland *et al*., 2011). Balota and Chaves (2011) and Zhou *et al*. (2017) and have reported great variation in microbial population with respect to legumes. Dhiman (2018) have also reported significant variations in the rhizobial population isolated from root nodules of *Dalbergia sissoo* seedlings, from different seed sources of Himachal Pradesh and Uttrakhand, grown under net house conditions.

### Authentication of rhizobial isolates associated with *albizia procera* root nodules

A total of 66 isolates 38 from Himachal Pradesh and 28 from Uttrakhand were isolated and accordingly different code was assigned to each isolate (Table 3) which is used in the further study.

These 66 isolates were screened for different authentication tests for rhizobia *viz*., Congo red test, Bromothymol blue test, growth in Hofer’s alkaline broth, ketolactose medium, glucose peptone agar and plant infection test (Figure1). Thirty eight isolates out of 66 gave positive results for all the authentication tests. None of the isolate absorbed congo red dye on CRYEMA medium after incubation period for 3-5 days at 28±2°C indicating that the isolates were rhizobia. Fifty four isolates were acid producer. The plant infection test revealed that only 38 out of 66 isolates nodulated the *Albizia procera* seedlings after 75 days of inoculation and these isolates were tentatively confirmed as *Rhizobium* spp.

**Figure 1:**
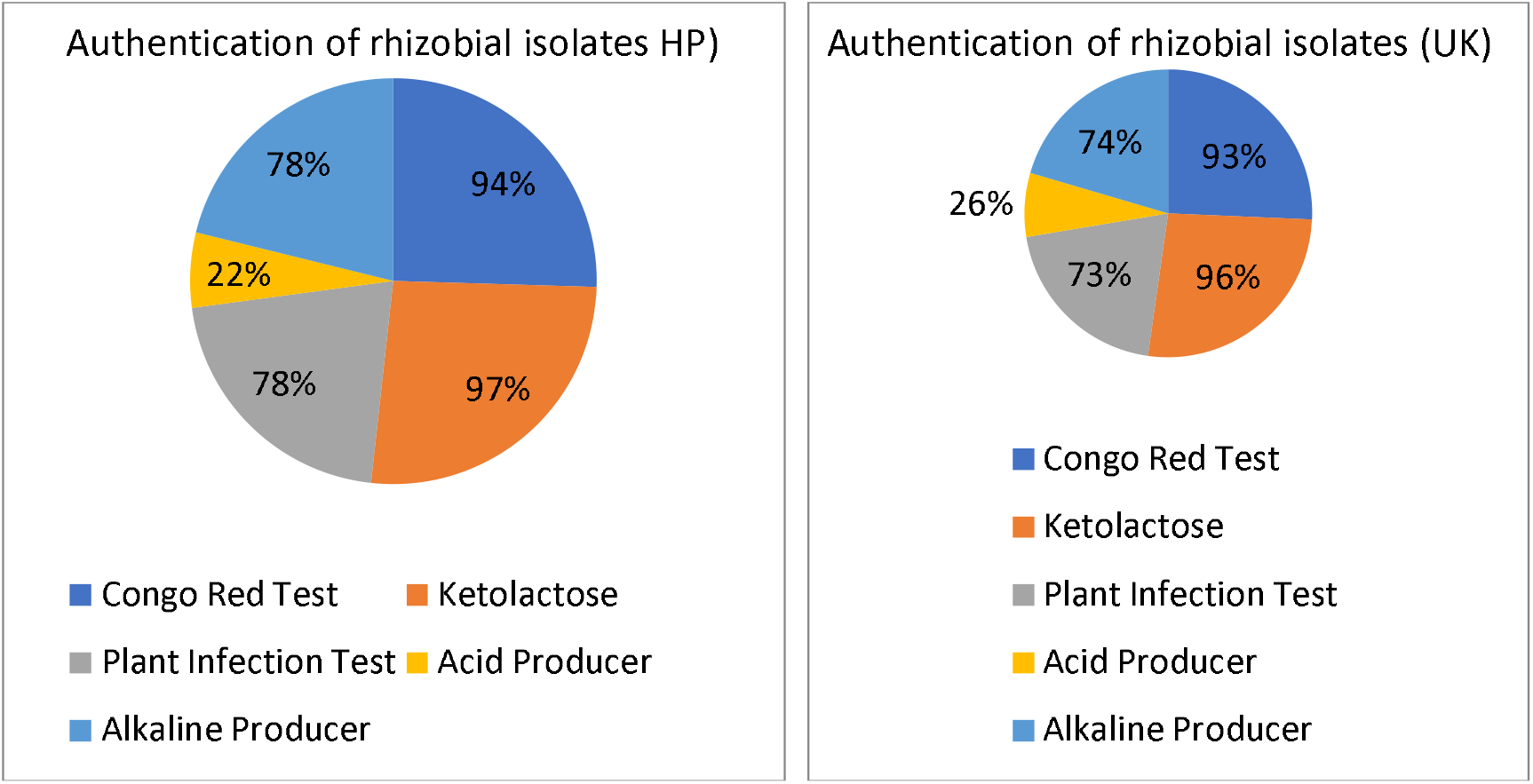
Authentication results of Rhizobial isolates isolated from *Albizia procera*.

Findings of the present investigations are in confirmation with Agarwal *et al*. (2012) who also reported that rhizobial isolates did not absorb red color when cultured in YEMA containing congo red dye, did not grow in Hofer’s alkaline broth and also did not utilize lactose and peptone. The results were further supported by Deshwal and Chubey (2014).

### Screening of selected rhizobial isolates for multifarious plant growth promoting traits

On the basis of different authentication tests, a total of 38 rhizobial isolates (nodulated the seedlings), out of which 21 isolates from Himachal Pradesh and 17 isolates from Uttrakhand were selected and further purified on YEMA medium. These isolates were screened for P-solubilization, siderophore, IAA and antagonistic activity against *Fusarium oxysporum, Fusarium graminearum, Pythium aphanidermatum, Alternaria, Rhizoctonia solani* and *Colletotrichum capsici*. The per centage of rhizobial isolates possessing P-solubilization, siderophore and HCN production among different locations as shown in Figure 2. The frequencies of rhizobia possessing PGP traits vary with respect to different locations. Among the total isolates from Himachal Pradesh 71.42 per cent were able to solubilize P, 76.19 per cent were able to synthesize siderophore on CAS medium and 47.61 per cent rhizobial isolate were HCN producers. Out of 17 rhizobial isolates from Uttrakhand, 82.35 per cent were P-solubilizers, 64.70 per cent were siderophore producers and 52.94 per cent were HCN produce (Figure2).

**Figure 2:**
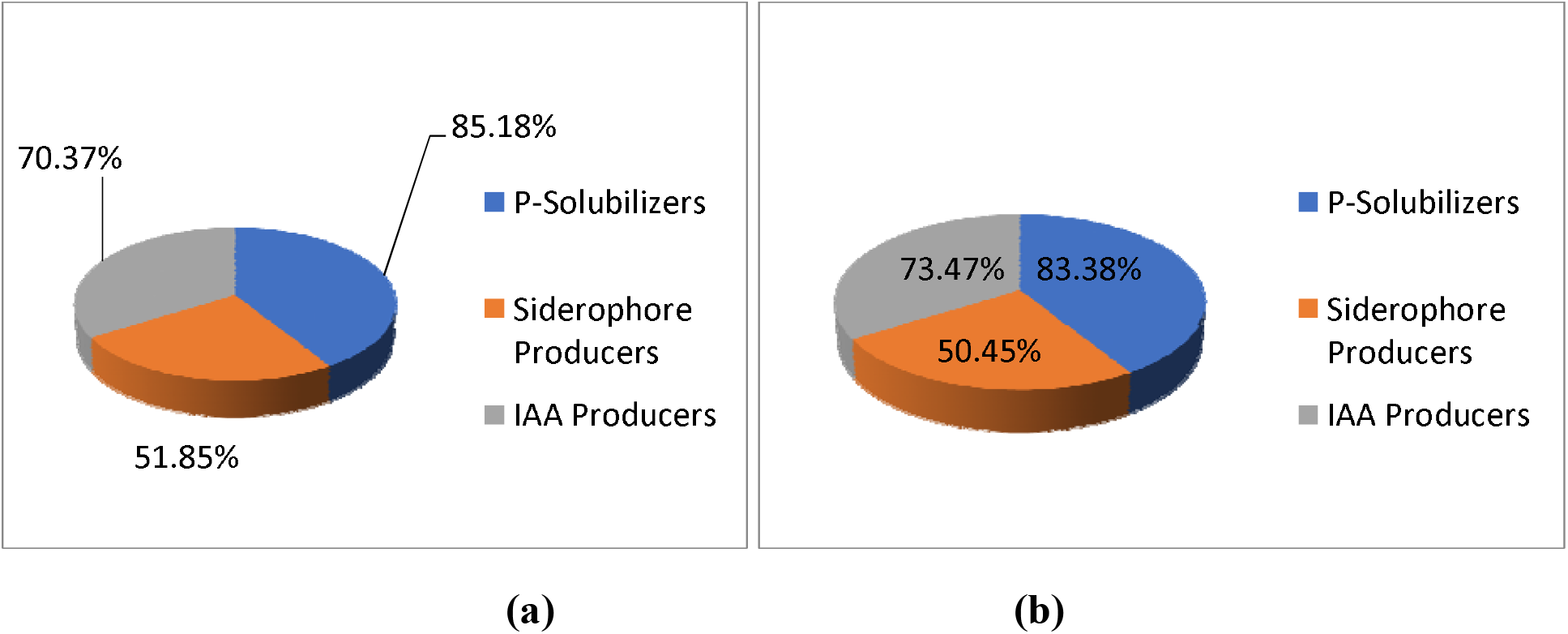
Percentage of Phosphorus solubilization, Siderophore production and IAA production by rhizobial isolates a) Himachal Pradesh b) Uttrakhand.

Rhizobial isolates which showed phosphate solubilization on PVK agar medium were further screened for quantitative estimation of P-solubilization in PVK broth (Plate 1).

**Plate 1:**
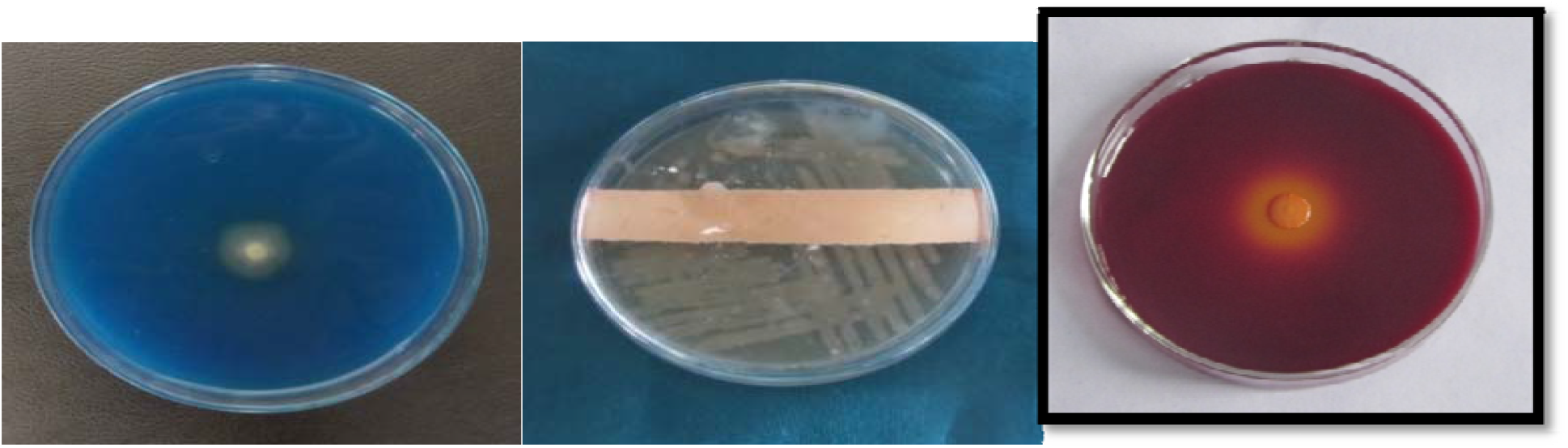
Screening of rhizobial isolates for various plant growth promoting traits of *Albizia procera*.

### Quantification of Plant Growth promoting Traits of selected rhizobial isolates

Legume nodulating bacteria (LNB), such as *Rhizobium, Bradyrhizobium* and *Mesorhizobium* have the ability to solubilize phosphorus. They increase availability of phosphorus to the plant by lowering soil pH through production of organic acids, chelation, ion exchange reactions (Delvasto *et al*., 2006) and mineralization of organic P by phospatases and lyases (Sharma *et al*. 2013). The results of the present study are in confirmation with Sridevi and Mallaiah (2009), Marra *et al*. (2012), Kumar and Ram (2014) and Rfaki *et al*. (2015) who reported P-solubilization by *Rhizobium* species.

Quantitative estimation of siderophore using CAS liquid assay of rhizobial isolates revealed that the maximum (106.30%) siderophore unit (SU) was observed in BA2 which wa statistically at par with JA4 (103.16%) while minimum (15.49%) siderophore production was observed by rhizobial isolate JA3. In case of rhizobial isolates from Uttrakhand maximum (108.12%) siderophore production was recorded with FA6 isolates which was statistically at par with siderophore production by rhizobial isolate IA5 (104.95%) as shown in Figure 3. The results are in line with Antoun *et al*. (1998); Duhan (2013) and Datta and Chakrabartty (2014) who also reported siderophore production by rhizobial strains.

**Figure 3:**
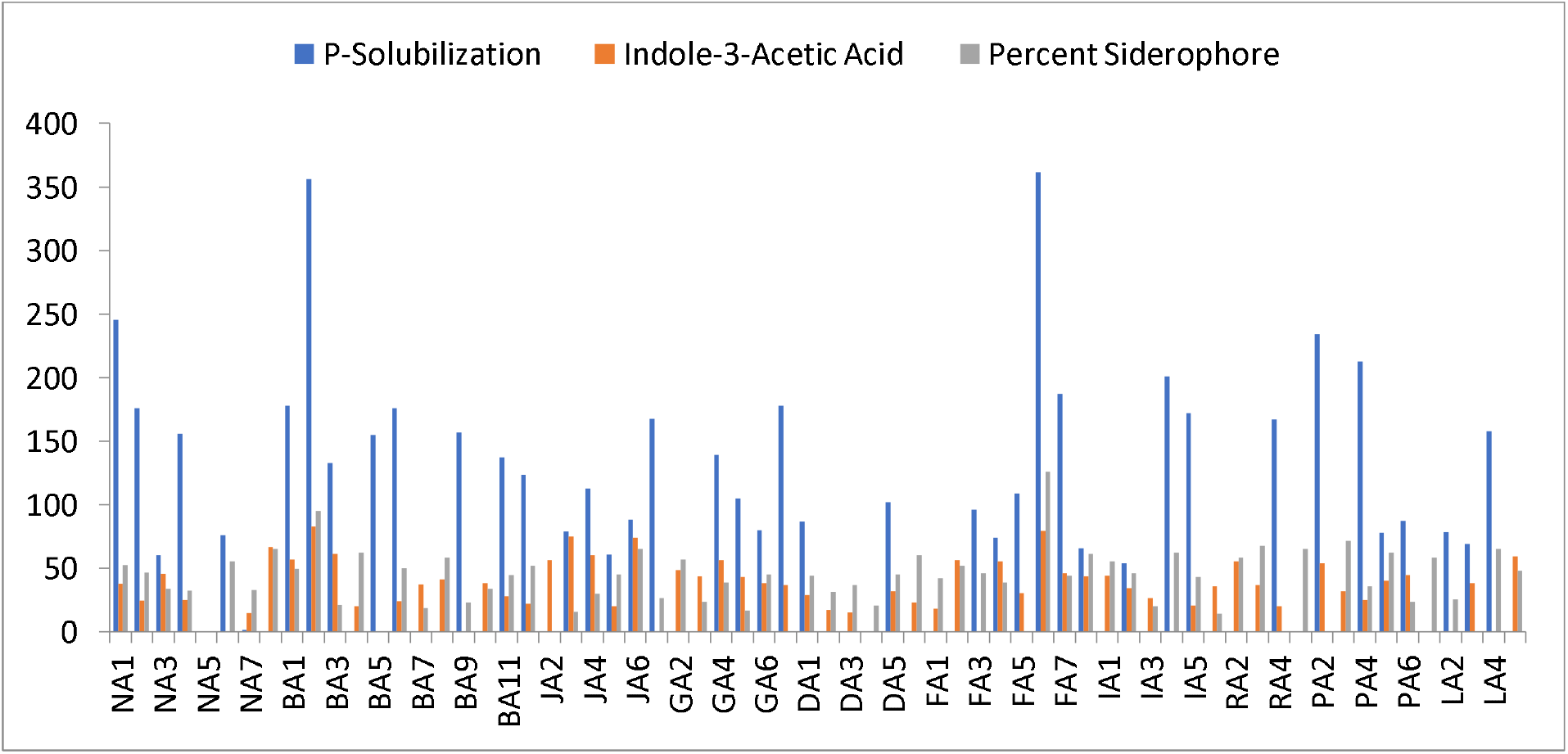
Plant growth promoting traits of rhizobial isolates from Himachal Pradesh and Uttrakhand.

The potential to produce siderophore by microorganisms improves the iron availability to plants (Wani *et al*., 2007; Ali and Vidhale, 2013; Kotasthane *et al*., 2017). Microbes release siderophore to scavenge iron from mineral phase by forming Fe^3+^ complex. Siderophore producing microorganisms protect plants at two levels i.e. by limiting growth of plant pathogens and triggering plants defensive mechanism (Ramos Solano *et al*., 2010). Dhull and Gera (2018) isolated 13 *Rhizobium* strains from the root nodules of cluster bean and reported that all the *Rhizobium* were able to produce iron chelating molecules.

IAA production by the rhizobial isolates varied from 14.95 to 85.52 μg/ml. Rhizobial isolate BA2 from Himachal produced maximum (82.75 μg/ml) amount of IAA from tryptophan (pH from 7.00 to 4.15) which was statistically at par with JA4 (79.19 μg/ml) whereas minimum (14.95 μg/ml) IAA production was recorded with rhizobial isolate NA7 after 72 hrs of incubation at 28±2°C. The rhizobial isolates from Uttrakhand, maximum (85.52 μg/ml) IAA production was recorded with FA6 isolate with reduction in pH from 7.00 to 4.21 and minimum (18.33 μg/ml) was in FA1 isolate with pH reduction from 7.00 to 5.85 (Figure 3). The maximum IAA production also coincides with maximum viable count.

Indole-3-acetic acid is a phytohormone which is known to be involved in root initiation, cell division, cell enlargement and resistance to stressful conditions etc. (Salisbury, 1994). Microbial synthesis of the indole acetic acid (IAA) stimulates and facilitates plant growth (Mohite, 2013). The results of present investigations are in conformity with Datta and Basu (2000) who reported high amount of IAA (99.7 μg/ml) production by *Rhizobium* isolated from root nodules of *Cajanus cajan* in L-Tryptophan supplemented basal medium. Sarayloo *et al*. (2014) have also reported Indole acetic acid (IAA) production in the range of 63.42 to 93.62 μg/ml by *Rhizobium rhizogenes*, isolated from soybean. Among all BA2 isolates from Himachal Pradesh and FA6 from Uttarakhand exhibited maximum plant growth promoting traits as shown in Figure 3.

### Antagonistic activity against fungal pathogens by selected rhizobial isolates

Plate 2 represents the per cent growth inhibition by selected rhizobial isolates against various fungal pathogens. *In vitro* antifungal activity of selected rhizobial isolates against phytopathogenic fungi including *Fusarium oxysporum, Fusarium graminearum, Pythium aphanidermatum, Alternaria, Rhizoctonia solani*, and *Colletotrichum capsici* were compared using dual culture method.

**Plate 2:**
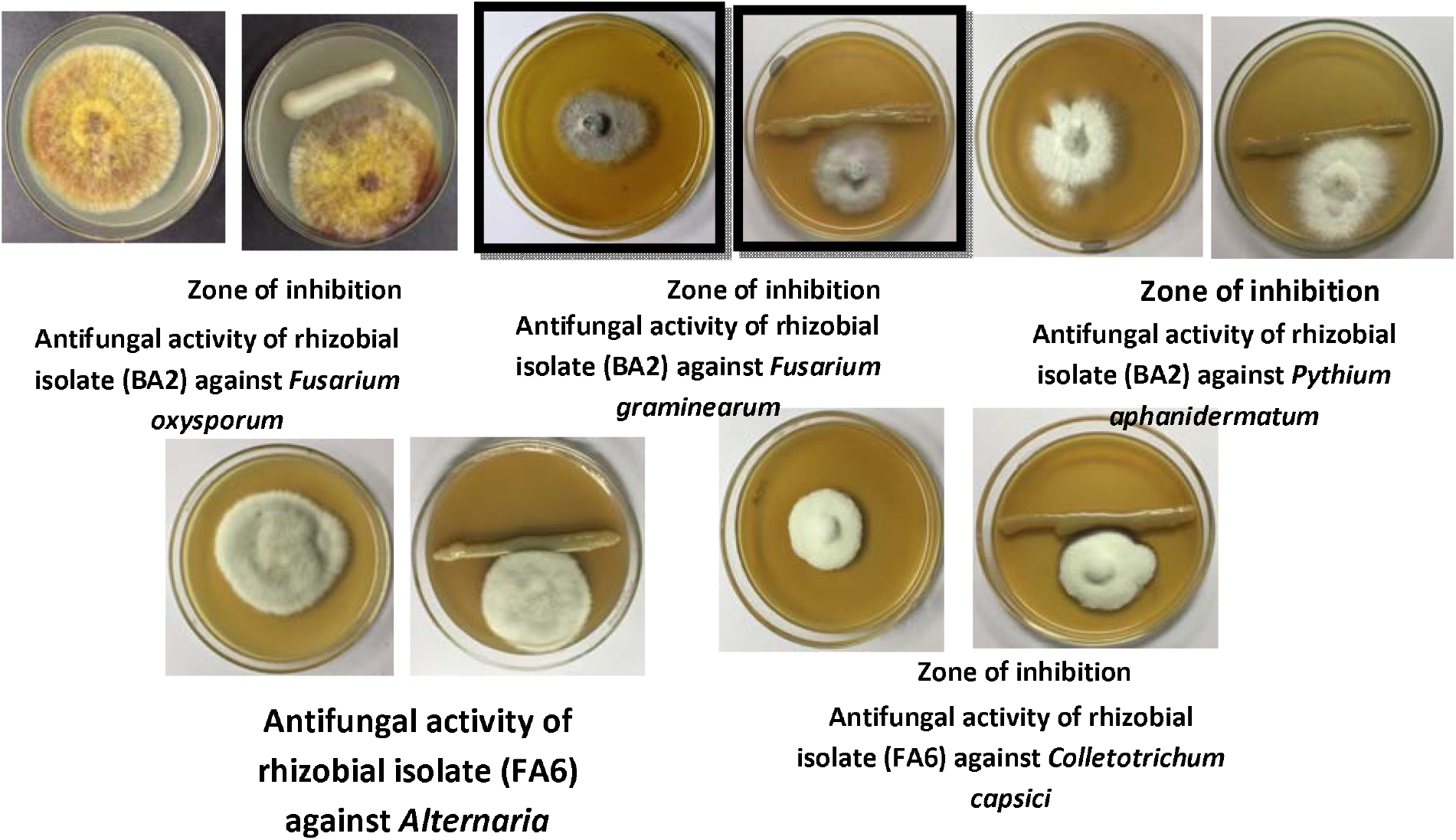
Antifungal activity of rhizobial isolate.

Rhizobia are also reported to induce systemic resistance and enhance expression of plant defense-related genes, which effectively immunize the plants against pathogens (Das *et al*., 2017). The formation of clear zone may be due to secretion of antifungal substance by the bacterial cell and that might have diffused in the medium indicating the inhibition of fungal growth.

Haque and Ghaffar (1993) also reported that *Rhizobium meliloti inhibited growth of M. phaseolina, R. solani* and *F. solani*. The results of our findings are also supported by Bardin *et. al*. (2004) who have reported *Rhizobium leguminosarum* bv. *viciae* strains R12 and R20, a PGPR for its ability to induce resistance in field pea against *Pythium*.

### Selection and evaluation of carrier material for survival of selected PGPR

Shelf life of selected rhizobial isolates viz., BA2, JA4, FA6 and IA5 with respect to different carrier materials i.e. charcoal, vermiculite and vermicompost was observed up to 6 months under laboratory conditions at 4^0^C temperature in refrigerator. The results showed that the density of initial rhizobial population differed from one carrier material to the other carrier material. The microbial analysis revealed that the selected rhizobial isolates (BA2, JA4, FA6 and IA5) retained desirable rhizobial population upto 4 months of storage at 4°C temperature and after that there was a decline in the population of rhizobia (Figure 4). For rhizobial strains BA2 and JA4, the initial highest viable count i.e. on 0th day was observed in Charcoal (165×10^8^cfu/g, 163×10^8^cfu/g) followed by vermiculite (161×10^8^cfu/g, 157×10^8^cfu/g). For FA6 and IA5 rhizobial strains the initial highest viable count i.e. on 0th day was also recorded in Charcoal (166×10^8^cfu/g, 162×10^8^cfu/g). It is also clear from the data presented in Figure 4 that viable count decreases with increase in number of days of storage. After 6 months of storage the maximum viable count for all the four rhizobial isolates was noticed in charcoal (118×10^8^cfu/g, 113×10^8^cfu/g, 115×10^8^cfu/g and 112×10^8^cfu/g). The carriers showed a different capacity for retention and survival of rhizobia. The viable count was significantly effective in charcoal in maintaining a higher viable population of rhizobia at initial stages. Rhizobial isolates BA2 from Himachal Pradesh and FA6 from Uttrakhand maintained the highest viable count in charcoal till 6 months of storage.

**Figure 4:**
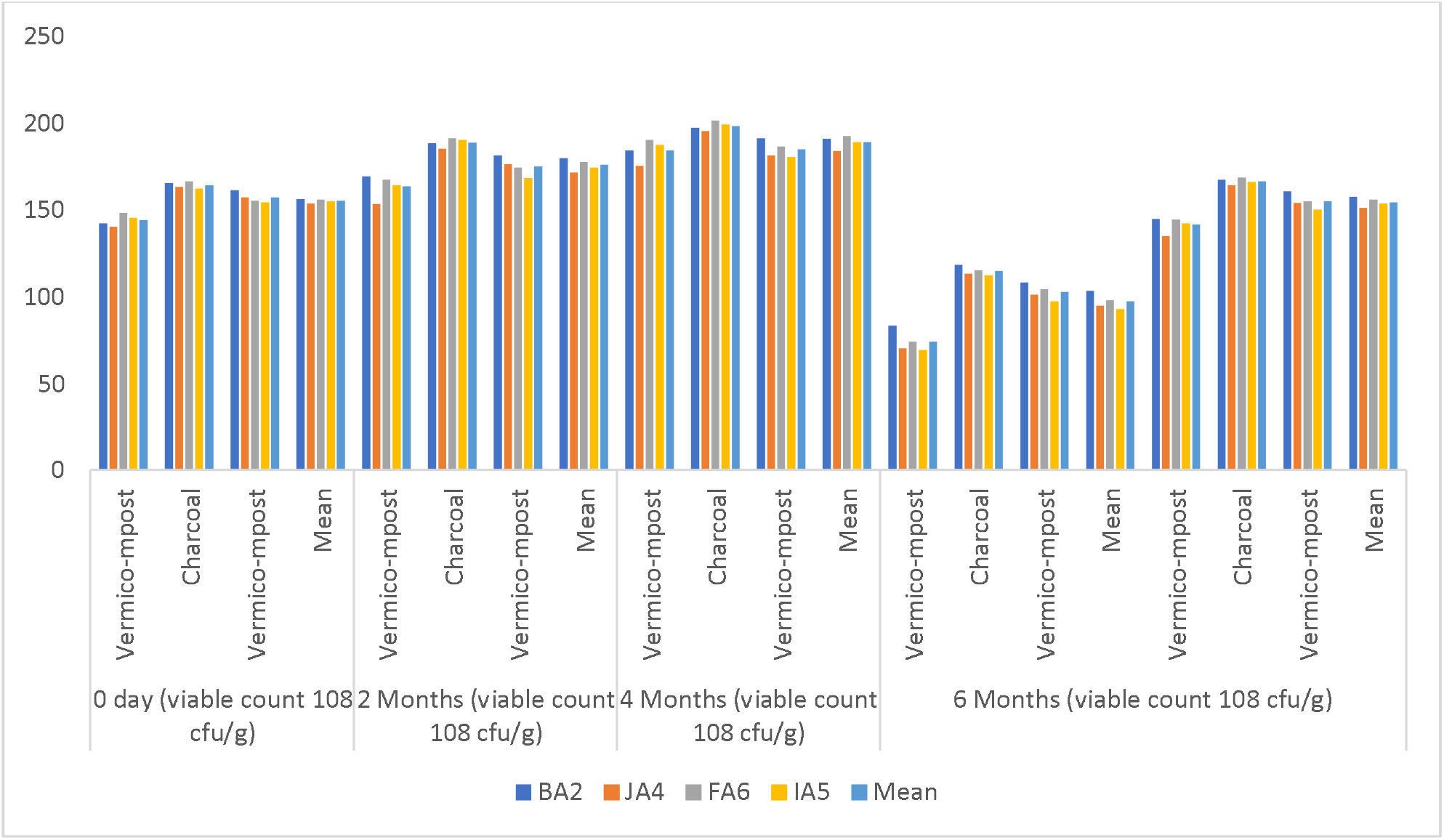
Shelf-life of rhizobial isolates on different carrier material.

The carrier is the delivery vehicle of living organism and it should deliver the right number of viable cells in good physiological conditions at right time, however, no universal carrier is presently available for PGPR inoculants. The shelf life of bacteria varies depending on the bacterial genera; carriers and their particle size. Carriers with smaller particle size have increased surface area, which increase resistance to desiccation of bacteria by increased coverage of bacterial cells (Dandurand *et al*., 1994). The increase in the viable count in charcoal as carrier material may be attributed to high carbon content, other sufficient nutrients, moisture and temperature during initial storage period. Deora and Singhal (2010) have also stated that the *Rhizobium* can be easily immobilized onto carriers like charcoal powder which can be applied as biofertilizer. Charcoal has widespread usage in agriculture (Antal and Gronli, 2003; Steiner *et al*., 2004b) and can easily be produced out of woody bio-mass. The properties of charcoal like inner large surface area, large pore size, maximum adsorptivity, good water holding capacity, non-toxicity, nutrient availability and its effectiveness to ameliorate adverse soil conditions make charcoal a good carrier for survival of rhizobia and bioinoculant production (Antal and Gronli, 2003; Glaser *et al*., 2002).

### Carrier based bioinoculant along with chemical fertilizer on growth and nitrogen fixation of selected seed source of *albizia procera*

Effect of carrier based rhizobial bioinoculant (prepared with charcoal) and chemical fertilizer (20 Kg/ha Nitrogen) on *Albizia procera* seedlings was studied under net house conditions. Seed coating as well as soil application of charcoal based rhizobial biofertilizer in general resulted in significant increase in various seedling parameters over control after six months of inoculation (Figure 5). In case of carrier based biofertilizer from Himachal Pradesh maximum shoot height (33.98 cm), shoot fresh weight (6.91 g), shoot dry weight (4.51 g), number of leaflets (85.34) and leaflet area (4.97 cm^2^) was observed in treatment T_2_ (Bioinoculant with BA2 strain). Similarly, significant increase in growth parameters of *Albizia procera* seedlings was observed when treated with rhizobial bioinoculant from Uttrakhand with treatment T_4_.

**Figure 5.**
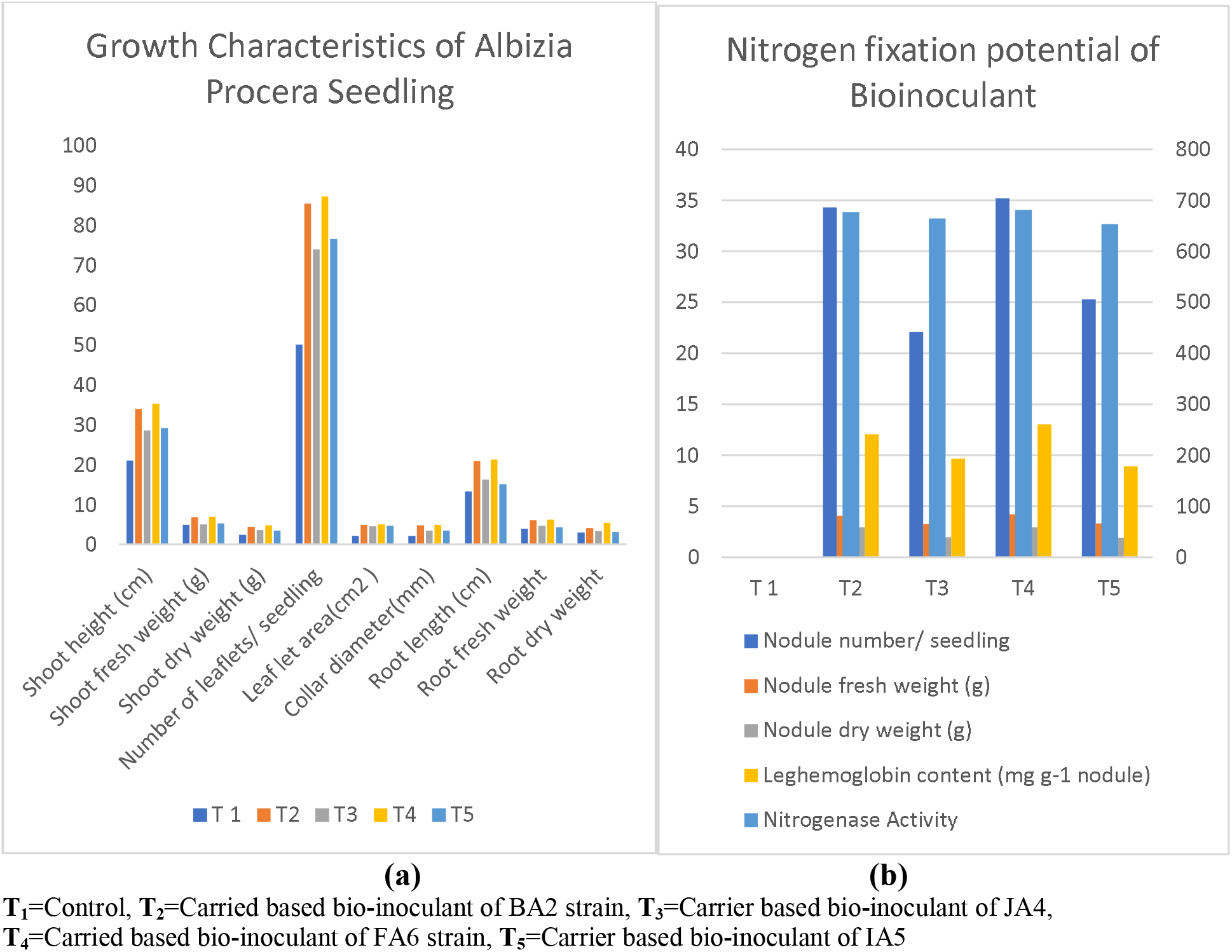
Effect of charcoal bioinoculant on growth and nitrogen fixation potential of *A. procera*.

Figure 5a revealed that root parameters were recorded maximum in seedlings under treatment T_2_ (Bio-inoculant with BA2 strain) for Himachal Pradesh and T_4_ for Uttrakhand.

The Nitrogen fixation potential of bioinoculant BA2 and FA6 were maximum in terms of maximum nodule number, nodule fresh weight, nodule dry weight, leghaemoglobin content and nitrogenase activity (Figure 5b).

Figure 6 showed that application of charcoal based rhizobial bioinoculant BA2 from Himachal Pradesh increased the shoot height upto 61.50 per cent and root length upto 58.45 per cent. Charcoal based application of rhizobial bioinoculant FA6 from Uttrakhand increased the shoot heightof *Albizia procera* seedlings upto 67.44 per cent and root length upto 60.72 per cent.

**Figure 6.**
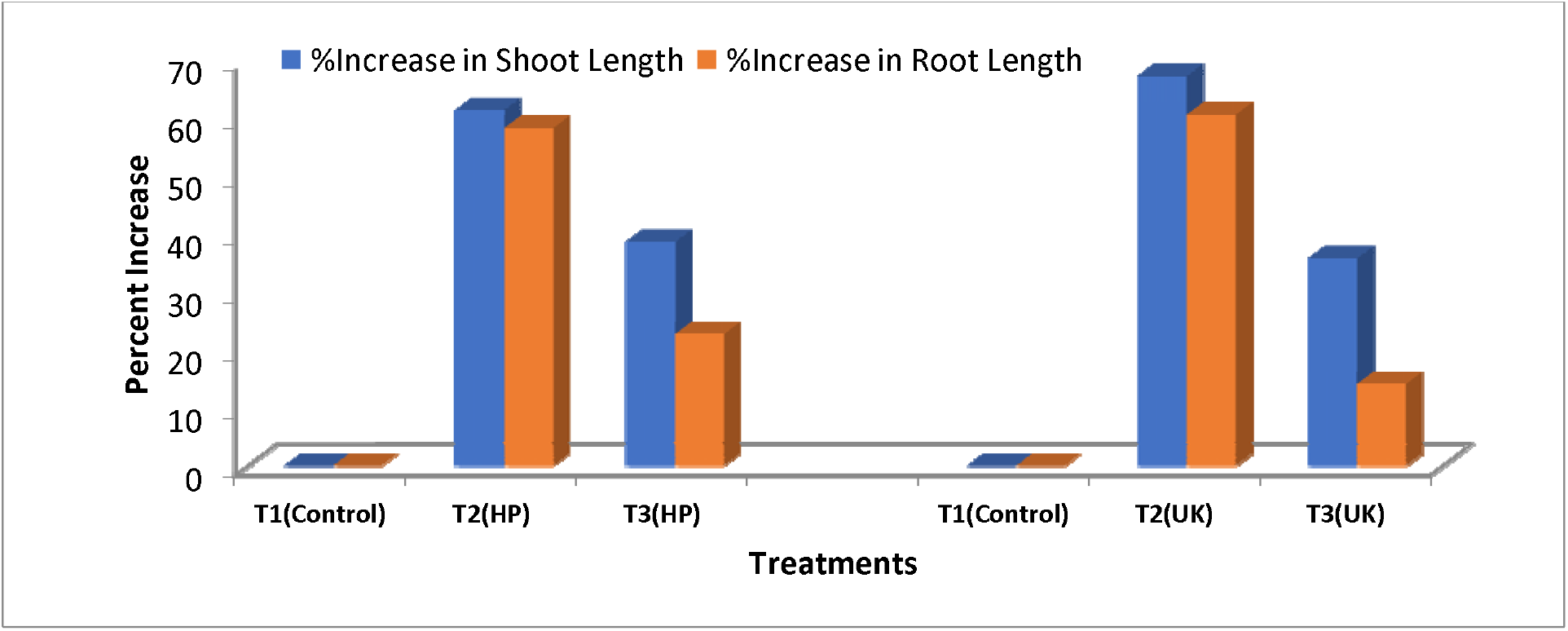
Per cent increase in shoot and root length after treatment with carrier based.

Among all the four charcoal based rhizobial bio-inoculants, the treatment T_4_ (prepared with rhizobial isolate FA6 of Uttrakhand) performed better for all of plant growth characteristics as compared to bio-inoculant prepared with other rhizobial isolates and uninoculated control (Figure 5& 6).

The effect of carrier based bioinoculant on growth and nitrogen fixation potential of *Albizia procera* was more pronounced as compared to liquid culture of biofertilizer. The charcoal based rhizobial inoculants have produced maximum growth and nodulation in *Albizia procera* seedlings. Our results for the present study are in agreement with Ravikumar (2012) who have concluded that the commercial inocula of *Rhizobium* with charcoal as a carrier used in seed inoculation of the pulses namely *Vigna mungo* (Linn.) Hepper. T-9 variety and *Vigna radiata* (Linn.) Wilzek S-8 variety has resulted in significant increases in growth and yield potential over the respective non inoculated control plants. Successful results by using commercial inocula of *Rhizobia* were obtained in field grown soybeans (Sharma and Tilak, 1974; Kapur *et al*., 1975; Dev and Tilak, 1976). Increase in the growth and yield of soyabean obtained due to rhizobial inoculation with charcoal might have been due to better physico chemical properties of charcoal as explained by Subba Rao and Tilak (1982), who obtained significant increase in grain yield in their test plants where they used charcoal as rhizobial carrier.

### Effect of carrier based bio-inoculant and nitrogen fertilizer on physico-chemical properties, nutrient availaibility and microbial count of soil

The data for physico-chemical properties of soil were recorded at the start and the termination of the experiments.

### Physico-chemical properties, nutrient status and microbial counts of soil (initial status)

The data pertaining to initial physico-chemical status of the soil is presented in the Table 4 revealed that the soil was nearly (pH 7.01), EC (0.34 dSm^-1^) was in normal range and organic carbon was 1.36 per cent. The available N (242.8 kg/ha), P (21.9 kg/ha) and K (274.4 kg/ha) contents were in medium range. The total rhizobial count was 50.9×10^6^ cfu/g soil on nutrient agar medium.

**Table 4.** Physico-chemical properties, nutrient status and total microbial count of soil mixture (initial status)

| PARAMETERS | VALUES |
| --- | --- |
| **1. PHYSICO-CHEMICAL PROPERTIES** |  |
| pH (1:2.5) | 7.01 |
| Electrical conductivity ($\text{dSm}^{-1}$) | 0.34 |
| Organic carbon (%) | 1.36 |
| **2. NUTRITIONAL STATUS** |  |
| Available Nutrients (kg/ha) |  |
| N | 242.8 |
| P | 21.9 |
| K | 274.4 |
| **3. TOTAL BACTERIAL COUNT ($\times 10^6 \text{ cfu g}^{-1} \text{ soil}$)** | 50.9 |

### Physico-chemical properties and nutrient status of soil at the termination of experiment

Figure 7 showed that application of charcoal based rhizobial bioinoculant BA2 from Himachal Pradesh increased the available N upto 33.15 per cent, P upto 77.94 per cent and K upto 44.27 per cent. Charcoal based application of rhizobial bio-inoculant FA6 from Uttrakhand increased the available N upto 34.96 per cent, P upto 78.84 per cent and K upto 44.85 per cent over initial status of soil.

**Figure 7.**
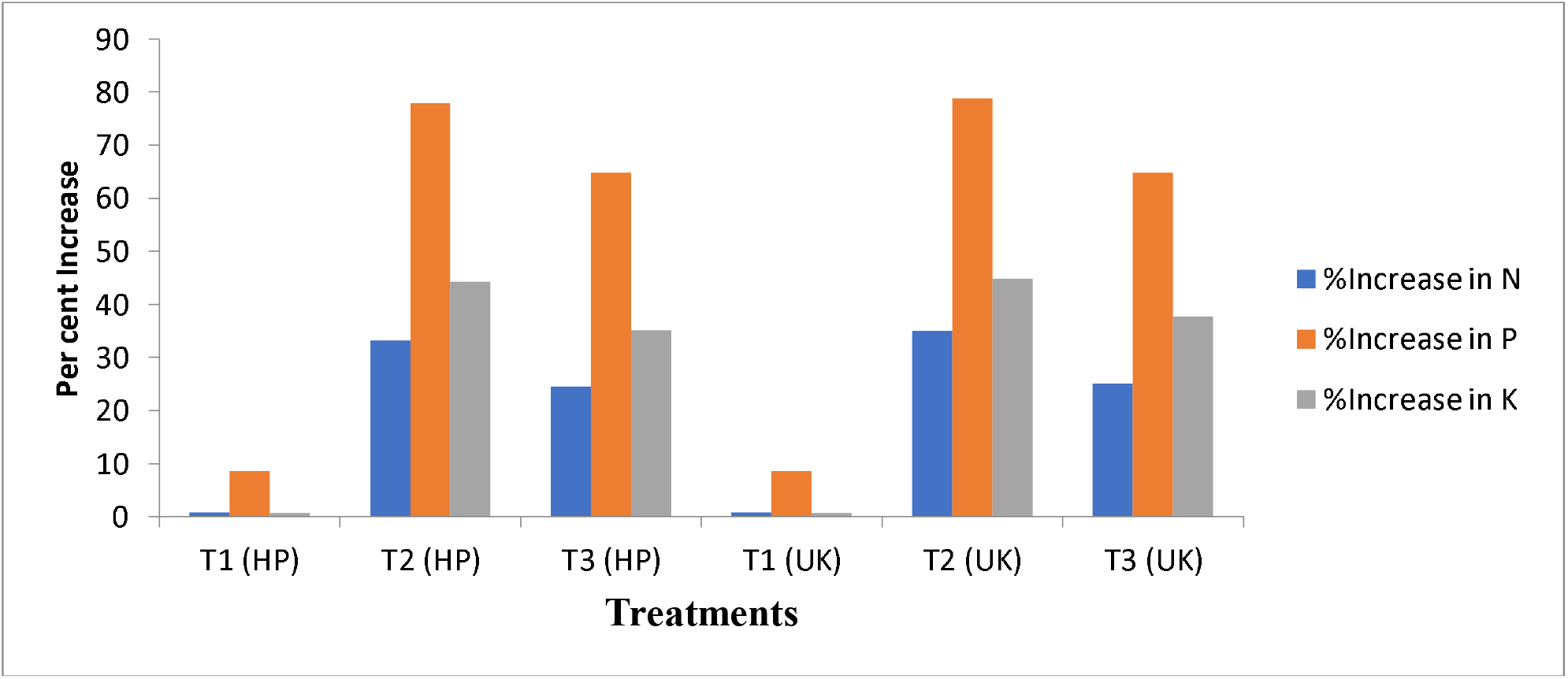
Per cent increase in available N, P, K after treatment with rhizobial bioinoculant over initial soil status.

The carrier-based formulations, in general, excel to register their effects on various plant growth parameters of *Albizia procera* over control. The increase in available N contents in soil may be due to mineralization of organic matter and fixation of atmospheric nitrogen by the symbiotic rhizobial isolates. These results are in conformity with that of Korir *et al*. (2017). Increase in the P availability might be due to the activities of phosphate solubilizing microorganisms, might have brought some P from unavailable pool to available pool. The solubilization of P in the rhizosphere is the most common mode of action implicated in PGPR that increase nutrient availability to host plants (Bhattacharyya and Jha, 2012).

### Effect of carrier based bioinoculant on soil enzymes (Dehydrogenase and Phosphatase

The conjoint application of rhizobial bioinoculant with chemical fertilizer significantly influenced the soil enzymes (Dehydrogenase and Phosphatase) over uninoculated control (Fig.8 (a&b)). In case of rhizobial bio-inoculant from Himachal Pradesh maximum (11.20 μg TPFg^-1^h^-1^) dehydrogenase activity was recorded with treatment T_2_ (Bio-inoculant with rhizobial strain BA2). In case of Uttrakhand bio-inoculant with rhizobial isolate FA6 (T_2_) also showed the maximum (11.39 μg TPFg^-1^h^-1^) dehydrogenase activity of soil which was greater than T_3_ and uninoculated control (T_1_). The maximum value for soil phosphatase activity (536.2 μmol L^-1^g^-1^h^-1^) was recorded under treatment T_2_ (Bio-inoculant with rhizobial strain BA2). However bioinoculant prepared with rhizobial strain FA6 (T_2_) from Uttrakhand also showed maximum (545.4 μmol L^-1^g^-1^h^-1^) soil phosphatase activity. The minimum activities for both the enzymes were recorded with uninoculated control (T_1_).

**Figure 8(a).**
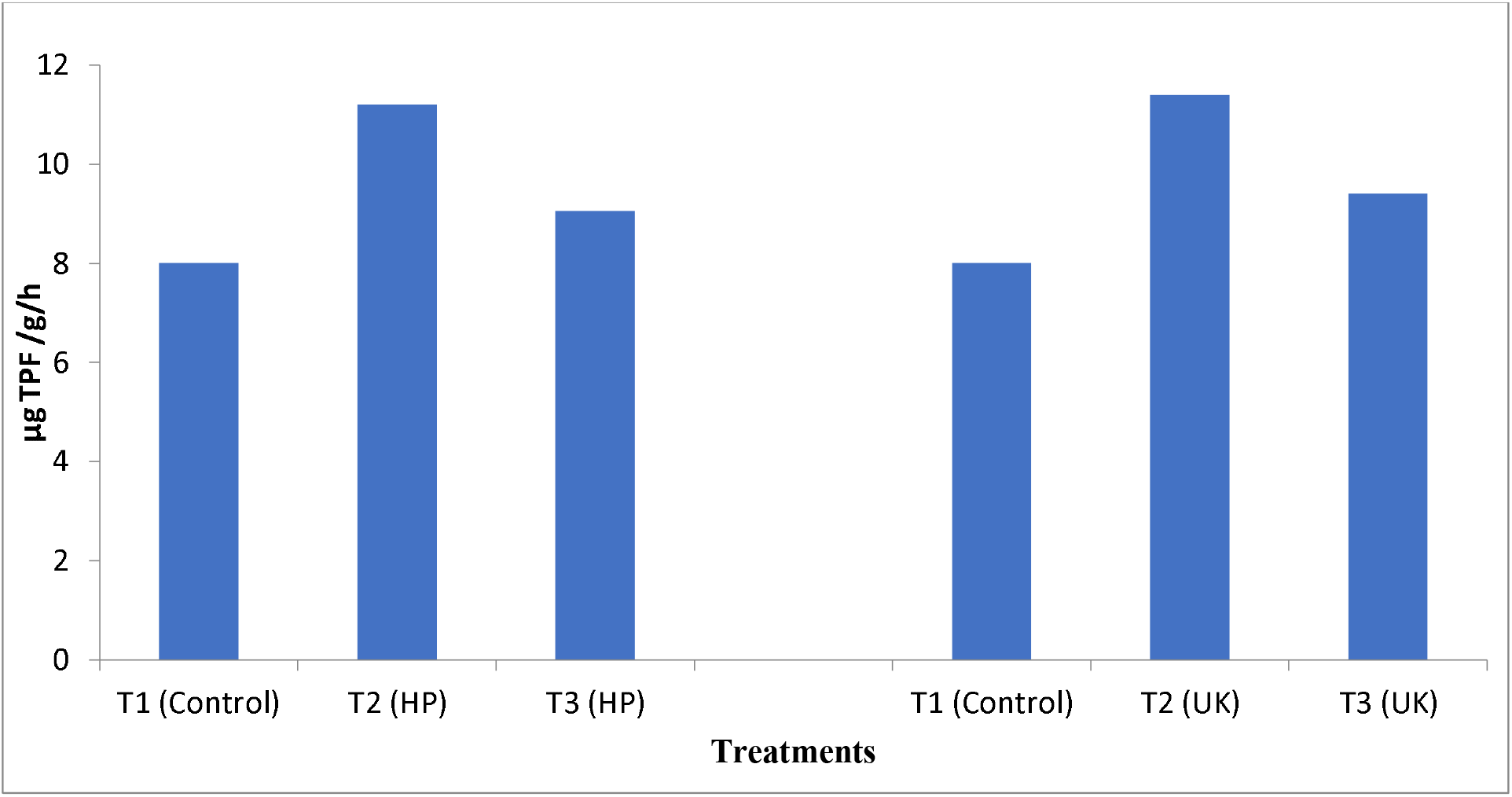
Effect of carrier based bioinoculant on soil enzymes (Dehydrogenase)

**Figure 8 (b).**
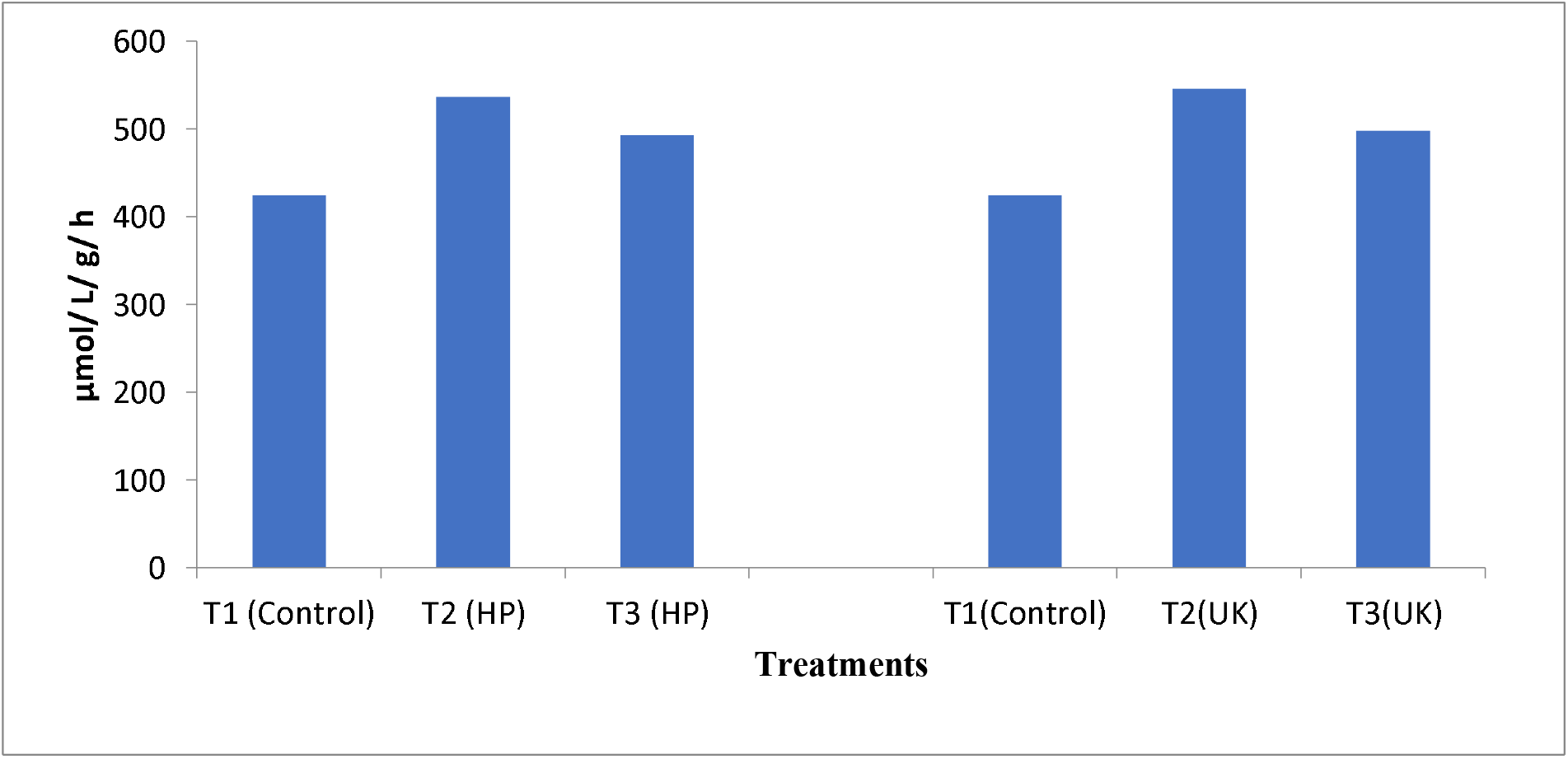
Effect of carrier based bioinoculant on soil enzymes (Phosphatase) Molecular characterization of selected rhizobial isolates.

Soil enzyme activities are incredibly important as they are direct linkage of soil microbial community to metabolic requirements and available nutrients present in soil. Higher enzyme activities in soil indicate the potential of soil to effect the biochemical transformations necessary for the maintenance of soil fertility (Rao *et al*. 1990). These findings are also in conformity with those of Mouradi *et al*. (2018) who also reported that inoculation of rhizobia together with chemical fertilizer increased the growth parameters (shoot height, shoot and root dry biomass) and soil enzyme activity.

On the basis of maximum plant growth promoting traits and results obtained from experiment shown by selected rhizobial isolates, two isolates i.e. one rhizobial isolate from Himachal (BA2) and one from Uttrakhand (FA6) were characterized by molecular identification. Molecular identification was done by sequencing part of the 16S rDNA. The amplification of the 16S rDNA was confirmed by agarose gel electrophoresis. The PCR products of both the rhizobial isolates were eluted and sequenced (Plate 3). The sequence data of the 16S rDNA of both the isolates was subjected to BLAST analysis. As 16S rDNA gene sequence provide accurate grouping of organism even at subspecies level it is considered as a powerful tool for the rapid identification of bacterial species (Jill and Clarridge, 2004). The sequence analysis of 16S rDNA of strain BA2 (Accession No.MK643828.1) and FA6 (Accession No.MK777968.1) showed maximum identity of 99.00 per cent to *Rhizobium* sp. The phylogenetic analysis of 16S rDNA sequence of the isolates along with the sequences retrieved from the NCBI was carried out with MEGA 6 using the neighbor joining method with 1,000 bootstrap replicates. The result of the phylogenetic analysis showed distinct clustering of the isolate BA2 with *Rhizobium leguminosarum* strain LMG 14904 (Accession No. NR 114989.1) and FA6 with *Rhizobium alamii* strain GBV016 (Accession No. NR 042687.1) (Figure 9). The results obtained are in line with Wang *et al*. (2006) who have isolated *Mesorhizobium* and *Rhizobium* originated from *Albizia* and *Acacia* sp.

**Figure 9.**
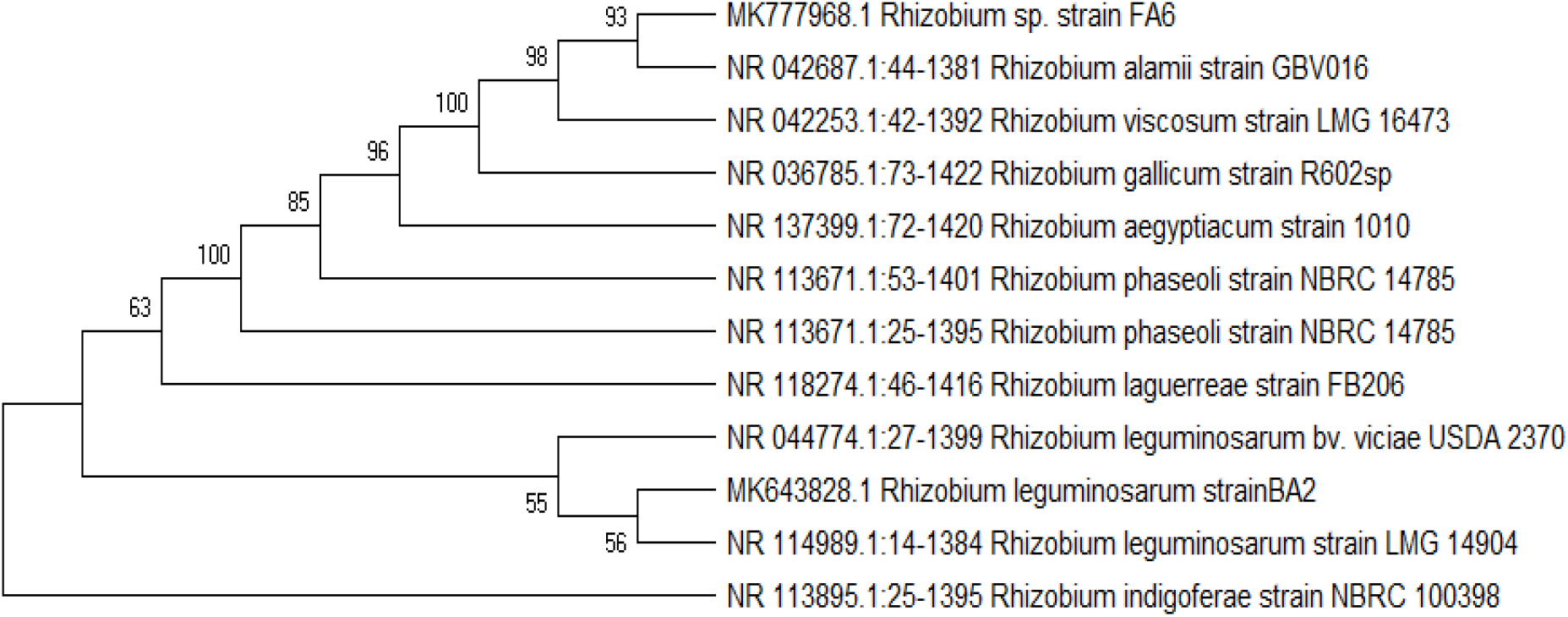
Neighbour joining tree based on 16S rDNA sequences showing the phylogenetic relationship of rhizobial isolates BA2 and FA6 with the analyzed sequences. (Accession numbers are mentioned within the phylogenetic tree corresponding to respective isolate)

## Conclusion

From the present investigation, it was concluded that selection of appropriate seed source with suitable rhizobial isolate can boost the growth and nitrogen fixation capacity of *Albizia procera* and play an important role in afforestation practices. On the basis of efficiency of selected isolates, the BA2 and FA6 have significantly increased shoot, root biomass and nitrogen fixation potential of *Albizia procera*. Hence, these isolates have enormous potential to be used as multifunctional biofertilizer for enhanced productivity of *Albizia procera* and to sustain soil health. The developed technology may be recommended to the farmers after field trials which can significantly reduce the dependence on chemical fertilizers coupled with eco-friendly approaches for production of quality seedlings of *Albizia procera*. It was also concluded that charcoal is the best carrier material for rhizobia as maximum viable rhizobial count was found in charcoal till six months of storage. Charcoal based bio-inoculant of rhizobial strains BA2 and FA6 alongwith 20 kg/ha nitrogen performed better for all the plant growth parameters. Thus, rhizobial strains BA2 and FA6 can be exploited as biofertilizer for improvement in *Albizia procera* seedlings. The developed technology may be recommended to the farmers after field trials which can significantly reduce the dependence on chemical fertilizers coupled with eco-friendly approaches for production of quality seedlings of *Albizia procera*.

## Acknowledgements

The authors are grateful to National Mission on Himalayan Studies (NMHS) and MoEF &CC for financial assistance

## Notes

### Competing Interest Statement

The authors have declared no competing interest.

